# How farming practices and livestock management affect Human-Wildlife Conflict intensity in Southern Ecuador: The case of the Spectacled Bear (*Tremarctos ornatus*) and feral dogs

**DOI:** 10.64898/2026.03.29.715147

**Authors:** Francisco Lopes, Mateo Peñaherrera-Aguirre, Rodrigo Cisneros

## Abstract

**Background:** Human–Wildlife Conflict is emerging as one of the most critical conservation and socio-economic challenges in the Ecuadorian Andes, where both rural livelihoods and native fauna are under increasing pressure. Small-scale livestock producers in the region depend almost entirely on a limited number of cattle, meaning that the loss of even a single animal can lead to severe economic hardship. In response, antagonistic actions against wildlife are frequent, further threatening vulnerable species. At the same time, the recent proliferation of feral dogs adds a new dimension to conflict, posing risks to both livestock and native fauna. Despite the growing severity of this conflict, little is known of its drivers, spatial patterns, and socio-ecological consequences. This study seeks to fill that gap by generating insights to inform targeted conservation strategies for community-based mitigation of conflict with spectacled bears and feral dogs.

**Methods:** To assess the drivers and dynamics of HWC in southern Ecuador, we conducted structured interviews with livestock owners, quantifying the frequency and intensity of conflicts across multiple species and evaluating whether farm composition and management practices predict conflict patterns.

**Results:** Our results reveal that large carnivores cause significantly higher economic losses than smaller predators; furthermore, feral dogs have emerged as the primary source of financial damage over the past five years. Farms with a greater proportion of forest edge were associated with a higher probability of severe conflict, particularly with large carnivores.

**Conclusions:** These findings underscore the urgent need for proactive strategies to promote coexistence. Identifying predictive variables of conflict risk is crucial for vulnerability assessments and the design of effective mitigation policies. Controlling feral dog populations is likely to be a critical step in safeguarding both rural human livelihoods and native biodiversity in the Andean landscape.

## Introduction

In recent decades, meat consumption has increased significantly, with estimates indicating a growth of approximately 58 percent, largely attributed to the rapid rise in global population (Alexandratos & Bruinsma, 2012; Chen et al., 2020). Demand for meat and milk is expected to continue increasing toward 2050 (Alexandratos & Bruinsma, 2012), driving the expansion of livestock production. This growth has been particularly strong in developing countries, where it is largely associated with the urbanization and population growth of these regions (Delgado, 2005). In contrast, production in the developed world, while remaining at high levels, has slowed or stagnated (Thornton, 2010).

Within the developing world, livestock inventories in Latin America have expanded at an especially rapid rate since the beginning of the twenty-first century, surpassing growth rates of the United States, one of the world’s largest livestock producers (FAO, 2024), as well as those in other regions (Williams & Anderson, 2019). South America alone accounts for about one quarter of the global cattle inventory (FAO, 2022), with Ecuador ranking among the countries with particularly high numbers of cattle (Baruselli et al., 2025), and therefore also undergoing an intense process of livestock expansion.

Although this increase in livestock production has generated economic benefits, such as contributing significantly to rising GDP per capita in Latin America (Williams & Anderson, 2019) and providing employment and livelihoods for millions of people, including poor farmers in developing countries (Thornton et al., 2006), it has also created major challenges. A central concern is that expansion of extensive cattle production frequently occurs at the expense of forests (Tyrell, 2019). It is estimated that 70 percent of deforested land in the Amazon is used as pasture (Williams & Anderson, 2019). Both deforestation and the increase in cattle production are associated with severe environmental consequences. Large-scale livestock production contributes to soil degradation and to alarmingly high levels of greenhouse gas emissions (Van der Hoek et al., 2016; Zhuang et al., 2019). Deforestation leads to extensive biodiversity loss (FAO, 2021), altered rainfall patterns, and increased temperatures (Pistora, 2024). It also reduces carbon storage capacity, thereby accelerating global warming (Li et al., 2022), while promoting soil erosion (Borrelli et al., 2013; Khodadadi et al., 2023) and water pollution (Kong et al., 2022). These effects are particularly concerning for cities such as Quito in the Ecuadorian Andes, which depend on páramo ecosystems for 85 percent of their water supply, a situation mirrored in many Andean cities (Buytaert et al., 2006).

A central issue linked to the conversion of native forests to pasture is the expansion of edge effects and the resulting intensification of Human–Wildlife Conflict (HWC). Edge effects arise where abrupt transitions form between ecosystems; in the Andes, deforestation to create low-complexity pastures fragments forests and produces additional forest–pasture boundaries (Murcia, 1995). Tropical species are particularly vulnerable to these effects due to their narrower temperature tolerances and limited dispersal capacities (Salisbury et al., 2012), making fragmentation more damaging than in temperate zones (Bregman et al., 2014). Edge expansion generates divergent species responses: some avoid edges, while others, including predators, exploit them to access livestock (Murcia, 1995). In Ecuador, where cattle frequently graze freely and near forests (MAE, 2019), this proximity increases predation risk. Simultaneously, expanding edges improve human access to native forests, facilitating hunting, logging, and retaliatory killings (Peres, 2001; Nyhus, 2016). Such persecution often stems from fear and misperception rather than verified losses (Goldstein et al., 2016; Albarracín & Aliaga-Rossel, 2018). Across Latin America, including Mexico, Costa Rica, Bolivia, Peru, and Brazil, cases of animal persecution and killing linked to such fears are well documented (Albarracín, 2010; Enríquez & Mikkla, 1997; Barnes, 1994; Catapani, Desbiez & Morsello, 2023). As deforestation proceeds, greater forest–pasture interfaces expose livestock to predation and predators to human retaliation, intensifying HWC and causing significant ecological and socio-economic harm (Conover, 2002; Treves & Karanth, 2003; Peterson et al., 2011). In the Ecuadorian context, Chastel (2018) reported that farms with a higher proportion of pasture directly bordering forested habitats, thereby creating greater edge zones, experienced significantly more predatory attacks than farms with less edge exposure. However, the study was limited by a relatively small sample size. More recently, Cisneros and colleagues (2025) analyzed the role of landscape composition and management variables in the Ecuadorian Andes and found that higher cattle losses were associated with greater distances between farms and farmers’ residences, infrequent farm visits, and larger pasture areas. Taken together, these findings suggest that variables related to farm composition, spatial configuration, and landscape matrix are likely to play a central role in shaping the intensity of HWC (Fig. 1).

**Figure 1.**
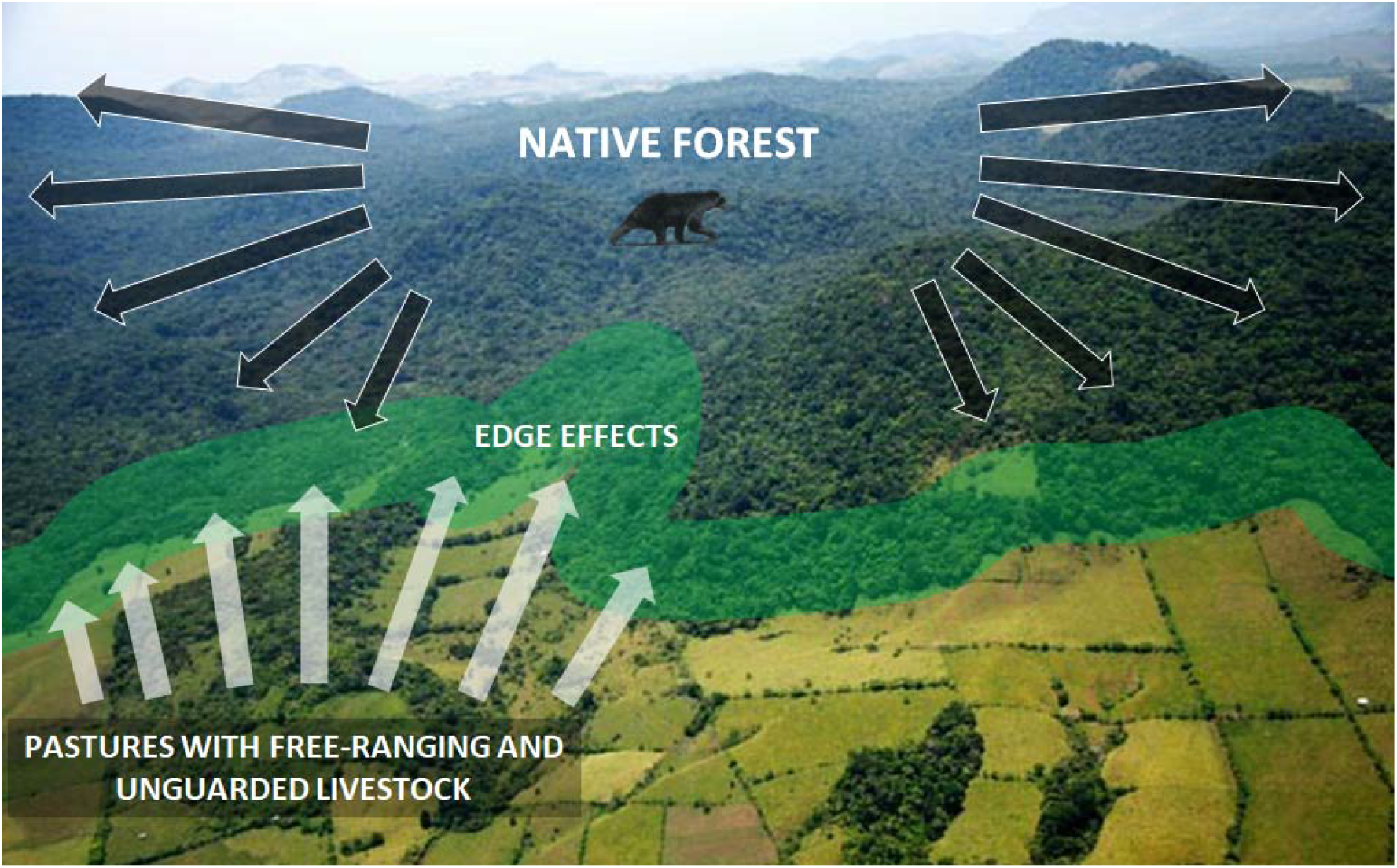
The functioning and growth of edge effects. Adapted from Cisneros et al., in press.

Although all carnivores are susceptible to Human–Wildlife Conflict (HWC) and retaliatory killing (Woodroffe et al., 2005), large-bodied species are particularly vulnerable due to their capacity to prey on high-value livestock such as cattle, posing substantial economic threats to rural livelihoods (Packer et al., 2005; Van Nieker, 2021; MAE, 2019). Human perceptions exacerbate this risk, as fear and “hyperawareness” of large carnivores amplify conflict likelihood regardless of actual threat levels (Treves, 2003; Dickman, 2010; Chapron et al., 2014). These dynamics are especially acute in areas of extreme poverty, where even minor livestock losses represent major financial shocks; in some cases, a single calf lost to predation may cost eight orders of magnitude more relative to income than in developed regions (Braczkowski et al., 2023). In Ecuador, where many communities rely heavily on cattle and face deep poverty, these socio-economic pressures intensify conflict, particularly with large carnivores such as bears, big cats, and canids (Cisneros et al., 2025; MAE, 2019). Collectively, these findings suggest that although HWC spans carnivore sizes, species capable of preying on cattle are disproportionately associated with severe conflict and retaliatory killing, underscoring the need for targeted, species-specific conservation strategies.

The spectacled bear (*Tremarctos ornatus*) is a keystone species in Andean ecosystems and widely regarded as an umbrella species due to its role in maintaining habitat structure and function (Crespo-Gascón & Guerrero-Casado, 2019; MAE, 2019; Vela-Vargas et al., 2021; García-Rangel, 2012). However, it is also a major contributor to Human–Wildlife Conflict (HWC) in Ecuador, where it preys almost exclusively on cattle unlike more generalist predators like pumas (*Puma concolor*) (MAE, 2019; García-Rangel, 2012; Cisneros et al., 2025; Narváez et al., 2022). Deforestation and the proliferation of free-ranging cattle in forest edge habitats amplify bear–livestock encounters, making the species a primary target for retaliation (MAE, 2019). Compounding this issue is the rapid rise of feral and free-ranging domestic dogs, particularly in areas lacking strong animal control policies (Home et al., 2017; Marshall et al., 2023). These dogs, often indistinguishable in behavior, breed freely, occupy forest edges, and have been documented killing livestock and competing with native fauna, including spectacled bears (Zapata-Ríos & Branch, 2016; Narváez et al., 2022).

Despite their growing impact, dogs are often overlooked in conflict assessments. Studies from Ecuador and beyond show that free-ranging dogs are responsible for many livestock deaths, yet native carnivores are frequently blamed due to negative perceptions (Home et al., 2017; Restrepo-Cardona et al., 2025). In Ecuador, dogs have been observed preying on cattle and harassing wildlife near protected areas, such as in the Madrigal del Podocarpus Private Reserve (Narváez et al., 2022; Restrepo-Cardona et al., 2025). Given their tendency to inhabit edge zones between human settlements and forests (Home et al., 2017), free-ranging dogs may be emerging as a key but underrecognized driver of HWC. Understanding their role is essential for accurately characterizing conflict dynamics and developing effective, evidence-based mitigation strategies that go beyond native carnivore management (MAE, 2019).

This study aims to evaluate the influence of farm composition and livestock quantity on the intensity of HWC in southern Ecuador. A further objective is to determine whether recently expanding populations of feral dogs are already present within the study areas and, if so, to assess the extent of their contribution to HWC as well as any factors that could inform future management strategies. We evaluate several hypotheses. [1] The first posits that the ratio of forest to pasture within each farm significantly influences conflict intensity, as this ratio reflects the extent of edge habitat and therefore the degree to which edge effects are experienced at the farm level. [2] The second hypothesis states that conflict involving large carnivores is expected to be more intense than conflict involving smaller carnivores, given that large carnivores typically cause greater economic losses (Woodroffe et al., 2005), which are among the most well-documented drivers of HWC. [3] The third hypothesis suggests that both the spectacled bear (*Tremarctus ornatus*) and feral dogs are likely to account for the highest levels of conflict among all species examined. [4] The fourth hypothesis proposes that feral dog populations are becoming more common in the region, and that their impact on livestock may already be comparable to, or approaching, that of the spectacled bear. [5] Finally, it is hypothesized that conflict involving large carnivores, notably the spectacled bear (*Tremarctus ornatus*), feral dogs (*Canis familiaris*), and the cougar (*Puma concolor*), will be significantly predicted by the extent of edge habitat within each farm and by the abundance of domestic animals, especially cattle and pigs. In contrast, the intensity of conflict involving small carnivores is expected to be primarily associated with the presence of poultry and guinea pigs, which are among the most traditional domestic species in rural Ecuador.

## Materials & Methods

### Study area and data collection

The study was conducted in various populated centers within the provinces of Zamora Chinchipe, Loja, and El Oro, located in the southern region of Ecuador.

Given the limited research on HWC and the absence of a public database documenting such events in this region, the study began with an exploratory phase to gather secondary information regarding reported HWC. Key informants, primarily officials from the Ministry of Environment, Water, and Ecological Transition of Ecuador, were consulted. Additionally, field researchers, and technicians working in the region across various fields of knowledge, who had heard from individuals affected by HWC, were also included as sources of information. Once affected individuals were identified, in situ visits were conducted, employing a chain sampling strategy (Newing, 2010) to carry out semi-structured interviews with farmers and individuals recommended by them. Local farmers from the communities were interviewed to investigate all aspects outlined in the study’s objectives. The sample included individuals who had experienced HWC as well as neighbours from the same area who had not experienced such conflicts in the last five years, ensuring a balanced representation of perspectives from both groups. Bushnell trap cameras were installed in 2024 in the Madrigal del Podocarpus Private Reserve to assess the presence of carnivores and feral dogs.

The altitudinal ranges represented in the sample spanned from 500 meters above sea level (masl) in locations such as Marcabelí, situated on the western foothills of the Andes, to dry valleys like Catamayo and Vilcabamba (1000 and 1570 masl, respectively). Rural areas corresponding to montane forests in Loja, exceeding 2200 masl, were also included, as well as the eastern foothills of the Andes in locations such as Yantzaza and Palanda (880 and 1120 masl, respectively). This diverse altitudinal representation ensured a comprehensive understanding of HWC across varying ecological zones within the study area.

We collected questionnaire data from 135 participants who own or rent farms and live in physical proximity to or within forest edge areas in the region. Interviews were conducted from December 2024 through March 2025. Only conflict events that have occurred within the last five years are considered in this study to ensure data standardization and facilitate meaningful comparisons. The ethical committee of the University of Porto approved this study. (Ref.: Proc. CE2025/p200).

### HWC intensity and frequency data

Species data for HWC intensity and frequency were not collected using an exploratory approach, meaning that interviewees were not asked to discuss any species with which they had experienced conflict freely. This decision was based on the fact that this specific step had already been conducted in the same region, the southern Ecuadorian Andes, by Cisneros and colleagues (2025). In that study, exploratory questions were used during interviews to obtain an initial list of all species causing conflict with local communities. The results provided a clear indication of which species are most vulnerable, and which cause the most significant damage to people. According to theory, species that present both high levels of conflict intensity and frequency are expected to be significantly more vulnerable to the effects of conflict (Treves & Karanth, 2003). Building on this knowledge, the present study focused on a more detailed analysis of conflict involving the eight most vulnerable species in the region, as identified by Cisneros and colleagues (2025) (Fig. 2).

**Figure 2.**
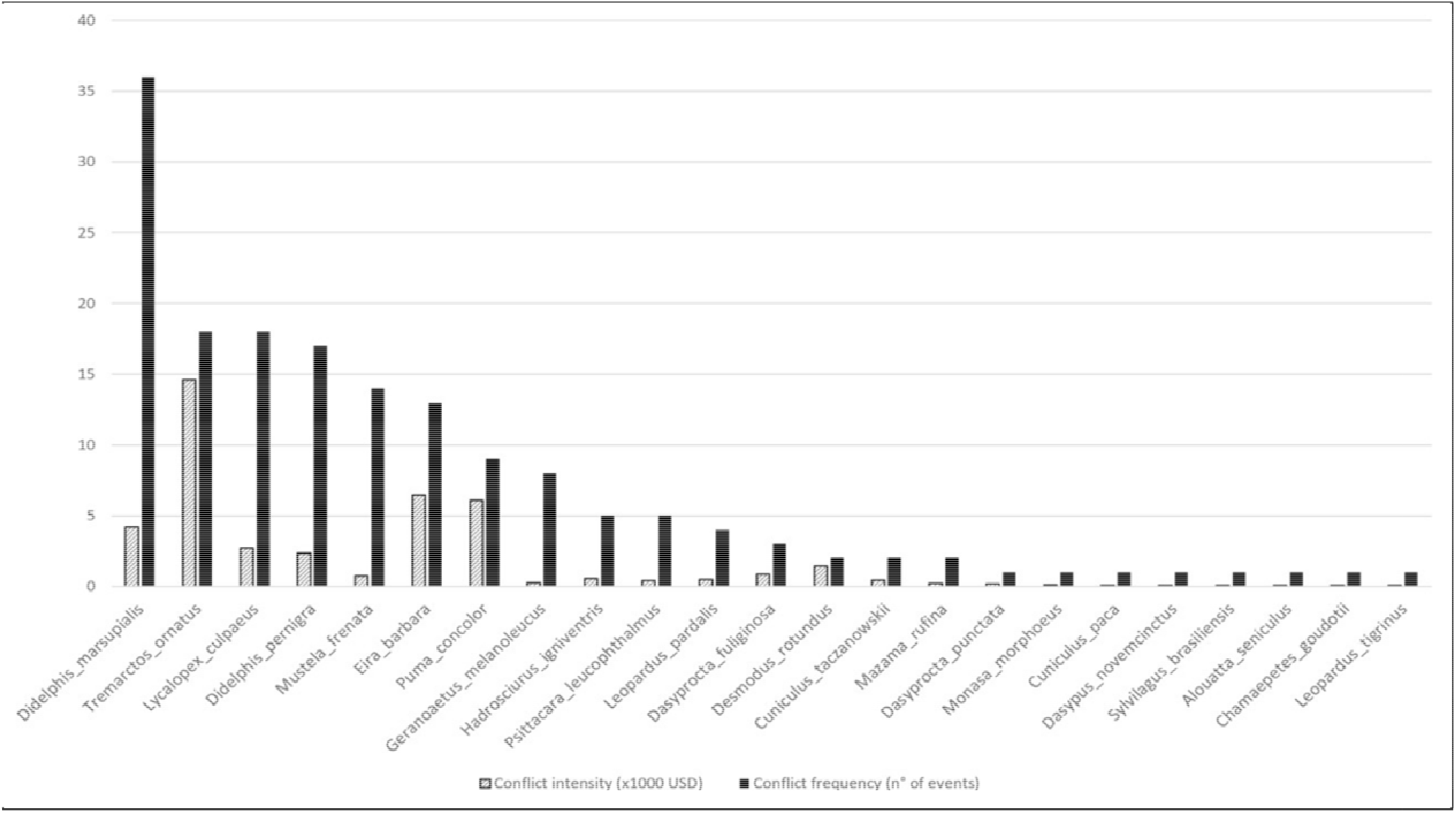
Intensity and frequency of conflict with species. Retrieved from Cisneros et al., in press.

Another reason for not repeating an exploratory methodology to identify species vulnerability to HWC, in addition to the statistically validated data from the previous study (Cisneros et al., 2025), is the challenge of accurately identifying species when working with local communities. People living in rural areas often confuse species, are uncertain about their appearance, or base their responses on exaggerated perceptions (Dickman, 2010). In the region, the initial exploratory interviews yielded reports of conflict involving species that do not occur on the South American continent, such as lions and even zebras (Cisneros et al., 2025).

For these reasons, the present study focused on the following species: the spectacled bear (*Tremarctos ornatus*), the cougar (*Puma concolor*), opossums (*Didelphis spp*.), the Andean fox (*Lycalopex culpaeus*), the long-tailed weasel (*Mustela frenata*), the tayra (*Eira barbara*), and the black-chested buzzard-eagle (*Geranoaetus melanoleucus*). Feral dogs (*Canis lupus familiaris*) were also included based on evidence from the literature. *Didelphis marsupialis* and *Didelphis pernigra* are almost indistinguishable to local people, which made it more practical to group them as *Didelphis spp*. Furthermore, all interviews were conducted using a high-quality printed sheet containing photographs of each species. This allowed interviewees to identify the species they were being asked about accurately and to specify which ones had caused conflict. Through this methodology, we aimed to minimize confusion, assumptions, and misperceptions, thereby ensuring that the results reflected reality as closely as possible.

This approach enabled the determination of specific values for the frequency of HWC, measured as the number of attacks caused by each species, as well as for the intensity of HWC, measured in terms of monetary losses in USD resulting from these attacks. The latter was used as the primary criterion variable in the statistical analysis.

### Statistical analysis

Several Hierarchical General Linear models were performed, examining the influence of a Pasture-Forest Extent factor (comprised of the standardized residuals of extension of the forest area per farm and the extension of the pastureland area per farm, controlling for the overall size of the farm), the number of animals kept per farm, and the statistical interactions between the latter predictors on the intensity of conflict with larger carnivorans and small carnivorans. Due to the high intensity of reported conflicts between farmers and spectacled bears, as well as cougars and feral dogs, subsequent examinations also considered the influence of cattle management and pig management on the intensity of these conflicts. Additionally, the effects of poultry management and guinea pig management were also considered as predictors of the intensity of conflict with small carnivorans. All analyses were performed using the car package in R version 4.5.1.

## Results

### HWC intensity and frequency for different species

From the 135 interviews conducted, the data largely align with the previously established hypotheses, although several unexpected patterns also emerged. As shown in Figure 3, and consistent with our predictions, the group of large carnivores exhibits a higher intensity of conflict compared with small carnivores. In other words, large carnivores are responsible for greater economic losses to rural households. This pattern persists despite small carnivores exhibiting a significantly higher frequency of attacks. This result is logical considering that, as discussed earlier, larger carnivores typically prey on more economically valuable livestock, particularly cattle, whereas small carnivores tend to attack smaller and less valuable domestic species.

**Figure 3.**
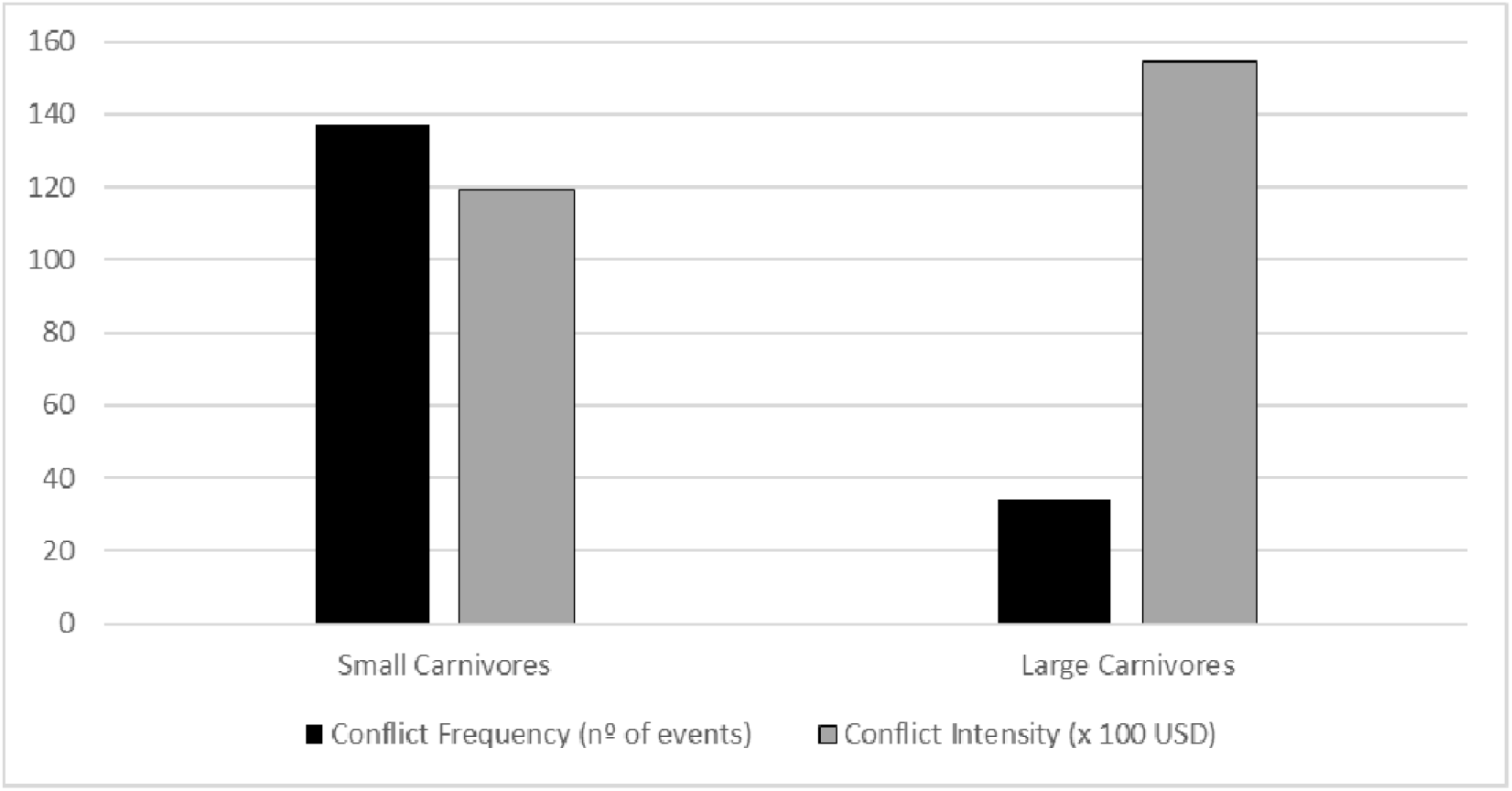
Differences in intensity and frequency of conflict between large and small carnivores.

When the data are examined at the species level (Figure 4), notable patterns become particularly evident. Over the past five years, feral dogs have represented the greatest source of economic losses for rural households. The second-highest losses are attributed to opossums (*Didelphis spp*.), a group classified as small carnivores. This finding raises an intriguing question. If, during approximately the same period, opossums caused more than twice as many attacks as any other species studied, and more than seven times as many as cougars (*Puma concolor*) or spectacled bears (*Tremarctos ornatus*), and ultimately caused the second-highest economic losses in the region, why are they largely absent from the literature on HWC and why are there so few reports of active persecution against them? This question will be addressed in greater detail in the discussion section.

**Figure 4.**
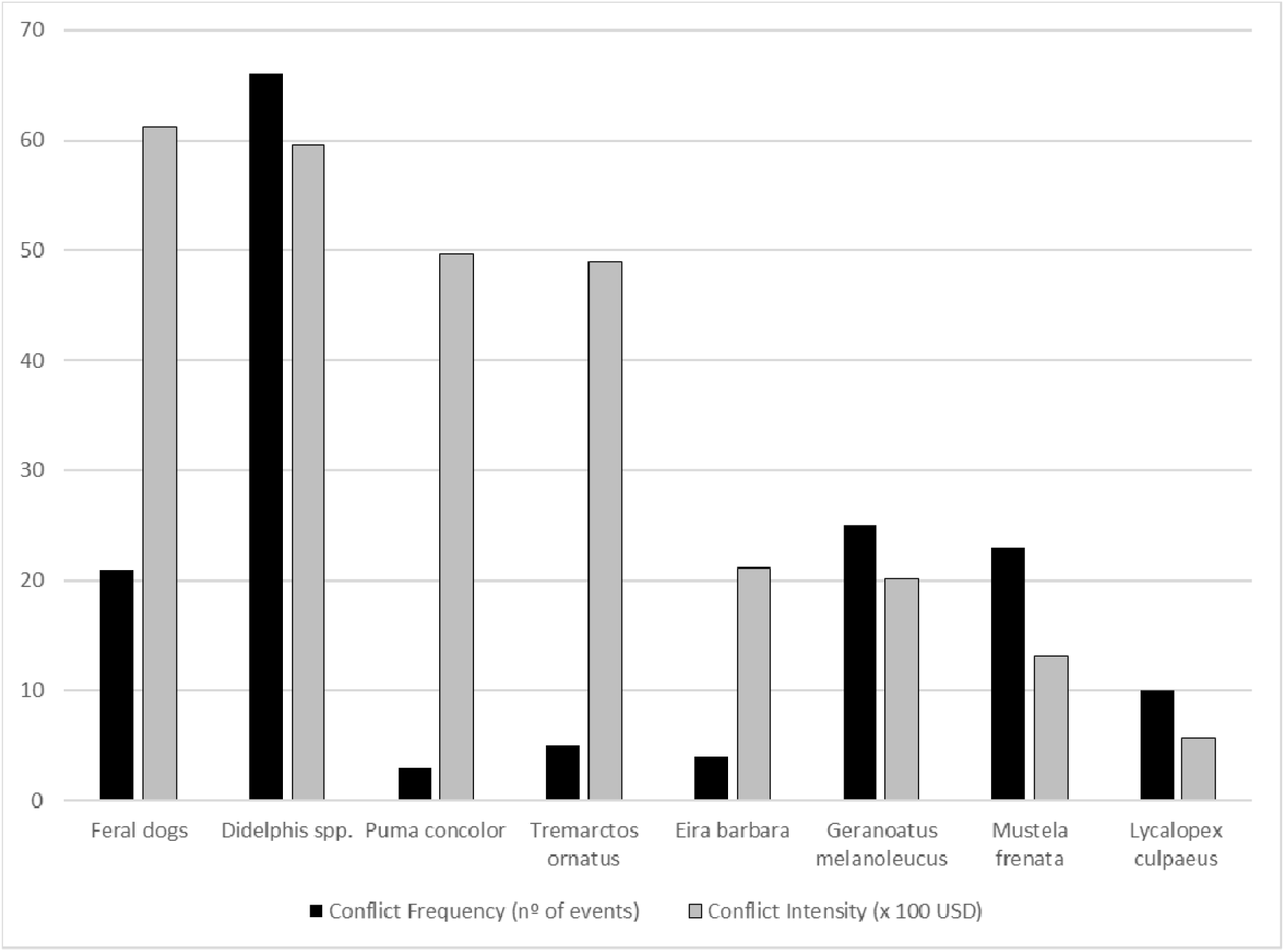
Differences in intensity and frequency of conflict between feral dogs, opossums, cougars, spectacled bears, tayras, black-chested buzzard-eagles, long-tailed weasels and Andean foxes.

Apart from the case of opossums, the remaining results are consistent with our initial hypotheses. Excluding opossums, the three species most responsible for higher intensity of conflict are the spectacled bear, the cougar, and feral dogs. When combined, these three species impose significantly greater economic pressure on rural households than all other carnivores studied. It is particularly noteworthy that both cougars and spectacled bears can cause substantial economic damage despite their relatively low frequency of attacks.

We also found that feral dogs are not only present in the study area but are already responsible for levels of damage comparable to, or even greater than, those caused by spectacled bears (Figure 5). Within the five years considered, feral dogs emerge as the single most significant source of livestock losses in the region. This finding underscores both the scale and the urgency of the problem, highlighting the need for feral dogs to receive greater attention in the broader HWC literature

**Figure 5.**
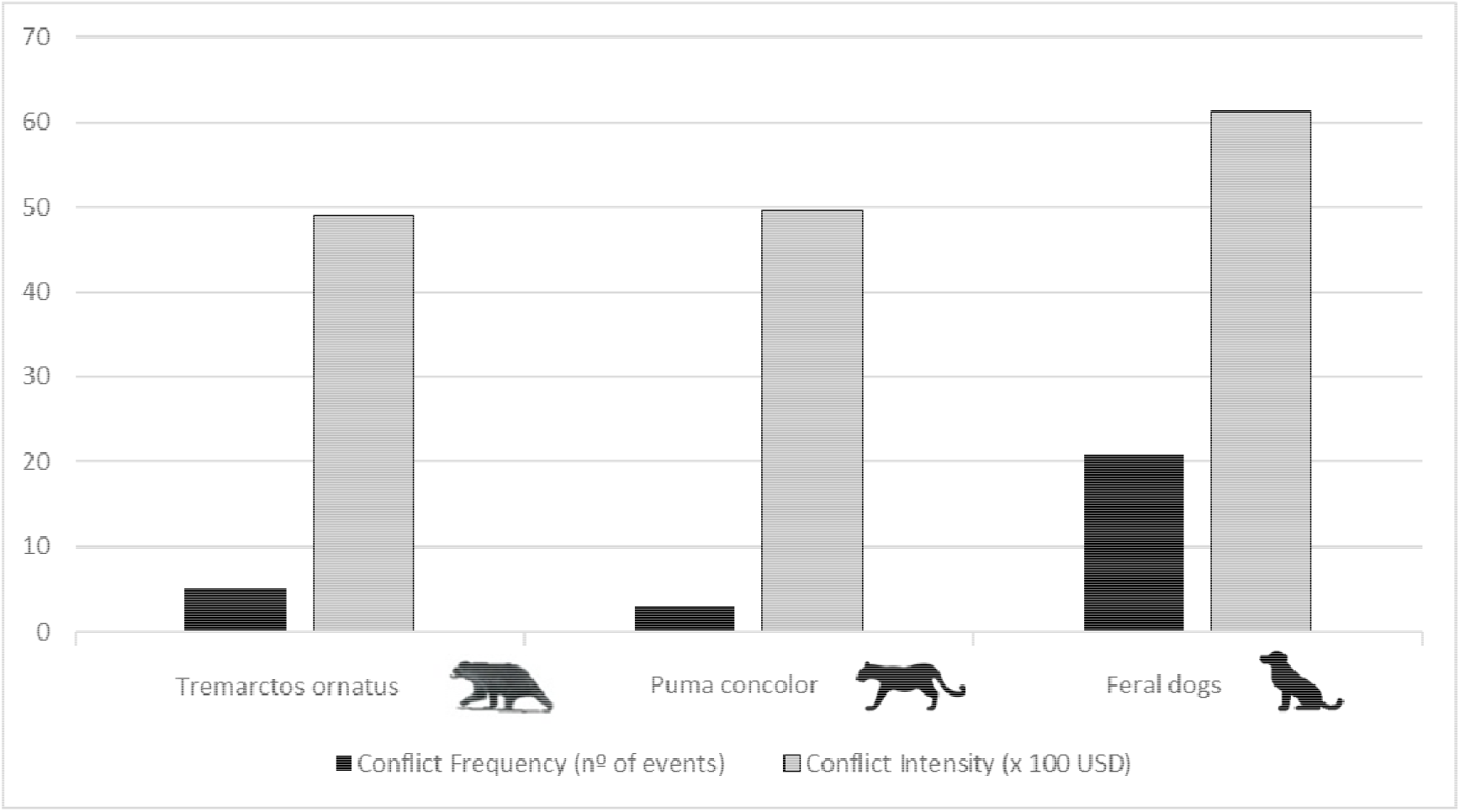
Differences in intensity and frequency of conflict between the large carnivores present in the study area.

Another noteworthy result concerns local perceptions of spectacled bear impacts. Previous studies have suggested that damage attributed to spectacled bears is often exaggerated due to cultural perceptions in Andean regions (MAE, 2019). The results presented here offer partial support for this idea. Spectacled bears are responsible for less economic damage than feral dogs and, notably, less than cougars. Interestingly, cougars, despite causing more economic losses than spectacled bears, appear to receive less negative attention, possibly because people have fewer direct encounters with them.

### HWC pathways

After residualizing the extension of pasture and the extension of forest for the terrain size, the residuals were standardized and exported for use in a unit-weighted factor estimation. The part-whole correlations between standardized residuals and a unit-weighted Pasture-Forest Extent factor indicated that this latent dimension loaded positively and significantly onto both the residuals for the extension of pasture and the extension of forest (r=0.996, p<.0001).

Subsequent hierarchical general linear models examined the main effects of the latter Pasture-Forest Extent factor and the number of animals kept per farm, in conjunction with their statistical interaction on the intensity of conflict with animals. The model examining the contribution of these predictors to the intensity of conflict with large carnivores reached statistical significance (p < .0001) and explained 30.8% of the variance. The analysis revealed that the Pasture-Forest Extent factor (β=0.284, p=.0009) and the number of animals kept per farm (β=0.489, p<.0001) positively predicted the intensity of conflict with large carnivores. In turn, the interaction between the Pasture-Forest Extent factor and the number of animals kept per farm did not reach statistical significance (β=−0.064, p=.7371).

Although the model examining the influence of the predictors mentioned above on the intensity of conflict with small carnivorans reached statistical significance (p = .0047), it accounted for only 9.4% of the variance. The analysis indicated that the Pasture-Forest Extent factor had no significant effect on the model (β = −0.024, p = 0.8441). Although the number of animals kept per farm had a significant effect, this value was small in magnitude (β = 0.164, p = 0.0060). Lastly, the model detected a negative and significant interaction between the Pasture-Forest Extent factor and the number of animals kept per farm (β = −0.523, p = 0.0182).

Similarly, the Hierarchical General Linear model, which included the Pasture-Forest Extent factor, poultry management, and guinea pig management as predictors of intensity of conflict with small carnivorans, reached statistical significance (p = 0.0042). However, the model explained only 9.6% of the variance. The analysis revealed that neither the Pasture-Forest Extent factor (b = 0.032, p = 0.8445) nor guinea pig management (b = −2.585, p = 0.0502) significantly predicted conflict with small carnivorans. However, poultry management had a positive and significant effect (b = 1.495, p = 0.0006) on the latter criterion variable.

Subsequent analysis examined the influence of the Pasture-Forest Extent factor, in addition to the main effects of farmers keeping cattle or pigs, on the intensity of conflict with specific types of medium to large carnivorans. For example, the Hierarchical General Linear model, with the intensity of conflict with spectacled bears as the criterion variable, reached statistical significance (p < .0001) and explained 34.6% of the variance. The model revealed that the Pasture-Forest Extent factor did not have a significant effect on the model, whereas both cattle (b = 0.330, p = 0.0005) and pig management (b = 2.332, p < 0.0001) had a positive influence on the intensity of conflict with spectacled bears. Similarly, the Hierarchical General Linear model, with the intensity of conflict with pumas, indicated the model was statistically significant (p < .0001) and explained 26.6% of the variance. The Pasture-Forest Extent factor (b = 0.150, p < 0.0001) and cattle management (b = 0.107, p = 0.0115) positively predicted the intensity of conflict with pumas. In contrast, pig management had no significant effect (β = 0.154, p = 0.1511) on the model. Lastly, the Hierarchical General Linear model with feral dogs as the criterion variable, reached statistical significance (p < .0001) and explained 29.6% of the variance. The analysis indicated that the Pasture-Forest Extent factor (b = 0.567, p < 0.0001), alongside cattle management (b = 1.029, p < 0.0001) and pig management (β = 1.140, p = 0.0043) had a positive and significant influence on the intensity of conflict with feral dogs.

### Trap camera documentation

To further corroborate the hypotheses already partially supported by the interview data, additional evidence was obtained through the use of camera traps placed at various locations along native forest edges within the study area. These camera traps provided crucial visual confirmation of the presence of feral dogs in these habitats. The devices recorded multiple instances of feral dogs, often appearing in groups, actively engaging in hunting behavior. In several of these images, feral dogs were observed pursuing the roost of a mountain paca (*Cuniculus taczanowskii*) (see Figures 6 and 7). This photographic evidence provides direct support for the conclusion that feral dogs are not only present in the landscape but are also actively interacting with native fauna, reinforcing the findings derived from interview data regarding their growing ecological and conflict-related significance.

**Figure 6.**
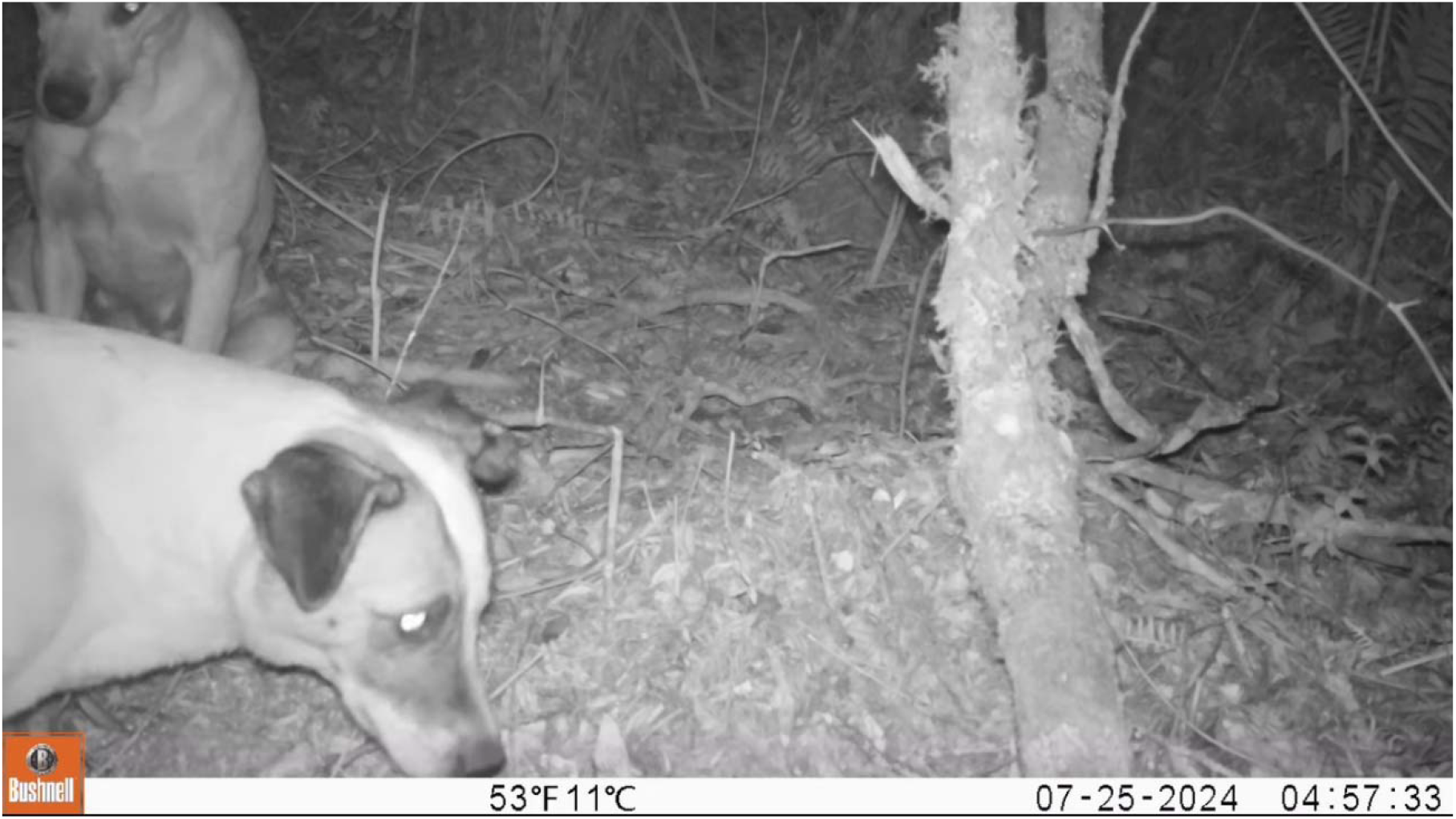
Two feral dogs harass a mountain paca (Cuniculus taczanowskii) roost, in Madrigaldel Podocarpus Private Reserve, 2024.

**Figure 7.**
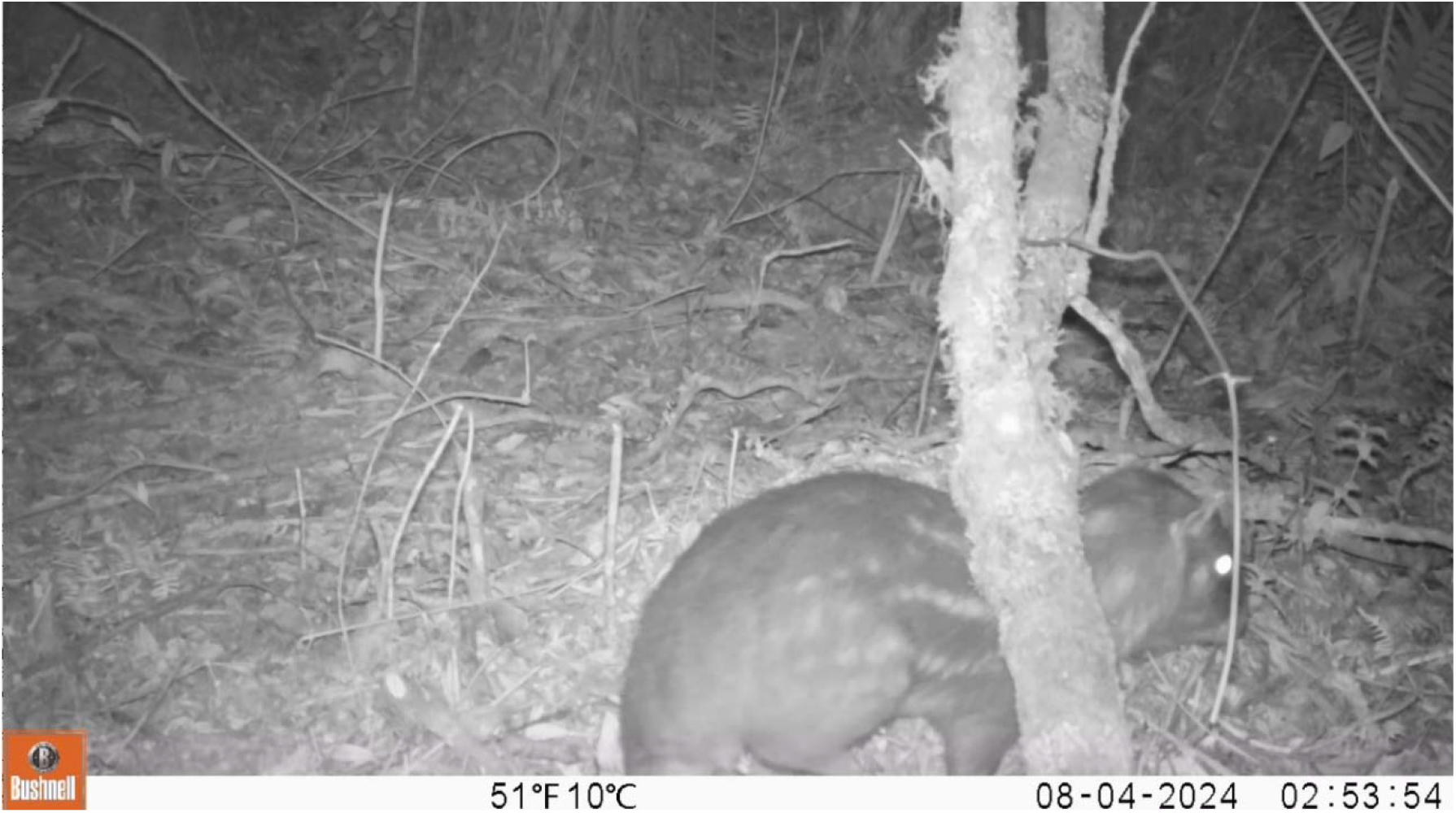
Mountain paca next to the roost in Madrigal del Podocarpus Private Reserve, 2024.

## Discussion

Differences in the intensity and frequency of HWC between species (Figs. 3–5) align with patterns reported in the literature. Large carnivores inflict greater economic losses on households than small carnivores, despite the latter attacking more frequently. This finding was expected, corroborated outside Ecuador (Woodroffe et al., 2005), and consistent with results from other regions of the country (Narváez et al., 2022; Cisneros et al., 2025).

Our findings extend this knowledge by highlighting the theory that may be driving these patterns. Extensive deforestation in Ecuador is converting forests into pastures (MAE, 2019), with approximately 60% of the population relying on livestock production for a living. However, most households subsist on fewer than a dozen cattle, which are often poorly managed, free-roaming, unguarded, and sometimes left unattended for weeks (García-Rangel, 2012; MAE, 2019; Cisneros et al., 2025). These herds graze in recently cleared and fragmented areas with reduced prey availability. Large carnivores, such as the spectacled bear (*Tremarctos ornatus*), which require extensive territories (García-Rangel, 2012; Vela-Vargas et al., 2021), are therefore compelled to prey on cattle. Evidence indicates that attacks are far less frequent in intact habitats with sufficient food, whereas fragmentation and edge effects substantially increase conflict (MAE, 2019).

Socioeconomic factors further intensify impacts. Rural livestock producers often live in extreme poverty (Braczkowski et al., 2023), with their household economies heavily dependent on a small number of animals (Cisneros et al., 2025). A single predation event can result in catastrophic losses, making large carnivores particularly vulnerable to retaliation and the primary source of conflict (Braczkowski et al., 2023). Small carnivores, in contrast, primarily prey on poultry (Williams et al., 2018; Amador-Alcalá et al., 2013; Narváez et al., 2022). Although these attacks are frequent and cumulative losses may be significant, they do not result in the sudden, catastrophic impacts characteristic of large-carnivore conflict.

Collectively, these ecological and socioeconomic factors explain the observed patterns of conflict intensity and frequency, as well as the species-specific differences illustrated in Figure 4. Two cases within species-specific results warrant further consideration. The first is the opossum (*Didelphis spp*.), which, over the past five years, has accounted for the highest frequency of attacks and the second-highest intensity, nearly matching the species responsible for the greatest economic losses. Despite this, opossums do not appear to generate the same vulnerability or antagonistic responses as large carnivores. Likely explanations include the gradual accumulation of impacts, primarily involving poultry, a resource of lower economic and cultural value than cattle (Amador-Alcalá et al., 2013; Cisneros et al., 2025), and reduced perceived threat, which diminishes retaliatory responses. The presence of a bear near a household evokes fear and insecurity in ways that smaller species, such as opossums, do not, likely prompting stronger retaliatory responses (Dickman, 2010; Prokop, Fancovicova, and Kubiatko, 2009). Ecological traits further mitigate vulnerability: opossums (*Didelphis spp*.) occur at higher densities, are more adaptable to human presence and effects, require smaller ranges, and are less affected by fragmentation (Greenspan et al., 2018; Magle et al., 2016; Veon et al., 2023). Consistent with this, the species is not listed as threatened in Ecuador (Tirira, 2021). Thus, while conflict with opossums (*Didelphis spp*.) is frequent and can be intense cumulatively, it is generally more tolerable and rarely catastrophic for households.

The second case is the cougar (*Puma concolor*). As a large carnivore, it might be expected to generate conflict levels similar to those of the spectacled bear (*Tremarctos ornatus*). However, despite ranking higher in conflict intensity than the bear, cougars appear to elicit less antagonism. This may be explained by their low frequency of attacks, which prevents them from being perceived as a persistent threat. Cougars are also less visible, spread predation across multiple domestic species (Narváez et al., 2022), and are more elusive (Arthurs et al., 2025), making retaliation more difficult. Together, these factors reduce the perception of risk and explain why, despite their measurable economic impact, cougars do not trigger the same level of conflict as spectacled bears.

### The growing threat of feral dogs

Although feral dogs are included in the results presented above, together with the remaining species, they require separate consideration. They represent one of the most novel and important contributions of this study, highlighting the scale and urgency of a problem that demands much greater attention in conflict research and management. The reviewed literature has already suggested that damage caused by feral dogs is increasing, particularly in the developing world (Lessa et al., 2016; Marshall et al., 2024; Home et al., 2017; Narváez et al., 2022). Based on this, we hypothesized that their contribution to conflict in Ecuador could rival that of the spectacled bear, traditionally considered the country’s most problematic species. The surprising and worrying results confirm not only that feral dogs have reached this level in the past five years, but that they have surpassed it. In the study region, feral dogs now represent the leading contributor to HWC in terms of both frequency and economic losses to rural households. This finding has profound implications. Uncontrolled domestic dogs are a widespread and largely unaddressed issue across the developing world. Populations increase without regulation or responsibility (Reece, 2005), and despite the scale of the problem, viable solutions remain scarce. If feral dogs are only now being studied but already cause substantial damage, the potential consequences in the coming years could be severe. Their impacts are multidimensional: they displace native carnivores (Zapata-Ríos & Branch, 2016), hunt and harass wildlife across taxonomic groups (Restrepo-Cardona et al., 2025), and increasingly prey on livestock (Narváez et al., 2022). Hunting in packs allows them to kill even large animals such as adult cattle, creating losses comparable to or greater than those caused by native carnivores (Home et al., 2017). If left unmanaged, this could escalate into a critical threat to ecosystems, native species, HWC dynamics, and the livelihoods of rural households.

The implications extend beyond the region where interviews were employed. While spectacled bears (*Tremarctos ornatus*) and other native carnivores will remain important and main contributors to conflict in many areas, feral dogs must now be recognized as a central driver of the problem. It is crucial to expand research on their distribution, impacts, and management, and to identify regions where the problem is emerging or intensifying. The context of this study, based in rural forest edge areas that have undergone partial urban development and lie close to national parks, may be especially conducive to feral dog expansion. Our camera-trap records confirm their presence along forest edges in the largest national park of southern Ecuador, and other reports indicate incursions inside protected areas in other regions of the country (Restrepo-Cardona et al., 2025). Together, these findings suggest that feral dogs may represent an emerging and underappreciated driver of conflict, with potentially far-reaching ecological and social consequences.

### Predicting Human-Wildlife Conflict Intensity

Identifying variables that heighten or reduce conflict intensity can guide future research, inform field methodologies, and help develop techniques to mitigate household economic losses, ultimately decreasing the motivation for retaliatory actions against wildlife. Our models offer promising insights into factors that influence conflict intensity. One of the most important variables tested was Pasture–Forest Extent, designed to capture the influence of edge ecosystems on the study area. This construct was calculated for each farm by assessing the relative proportions of pasture and forest, providing an approximate measure of the amount of edge habitat present within farm boundaries. While this measure has limitations, notably the exclusion of edge effects that occur beyond farm borders, it nonetheless yields valuable insights.

Conflict intensity with large carnivores, the most concerning type of HWC (Woodroffe et al., 2005; Packer et al., 2005; Van Nieker, 2021), was significantly predicted by Pasture–Forest Extent. Farms with greater edge habitat experienced higher conflict intensity, consistent with the literature linking deforestation and fragmentation to increased HWC (Murcia, 1995; Peres, 2001; MAE, 2019). This highlights the importance of identifying and mitigating edge effects resulting from the extensive conversion of forests to pastureland. Interestingly, the same factor was also associated with conflict involving pumas and feral dogs. This suggests that edge areas not only attract large native carnivores coping with fragmented habitats but may also facilitate the establishment of feral dog populations. Such areas provide both easy access to livestock and proximity to human settlements, from which feral dog populations originate (Murcia, 1995; Home et al., 2017).

Additional variables also provided valuable insights into the drivers of conflict intensity. For large carnivores overall, intensity was predicted by the number of animals per farm. This suggests that carnivores are more likely to generate severe events of conflict on farms with a larger number of animals, making it a practical indicator of vulnerability at the farm scale. For the species-specific model of the spectacled bear (*Tremarctos ornatus*), cougar (*Puma concolor*), and feral dogs, we replaced number of animals with the presence of specific domestic species to test whether certain livestock types were associated with higher conflict intensity. The results were consistent and informative. Conflict with spectacled bears was most strongly associated with the presence of cattle, followed by pigs, which reflects both the literature and local reports indicating that bears show a marked preference for attacks on cattle (Narváez et al., 2022). The presence of cattle also predicted the intensity of cougar conflict. For feral dogs, both cattle and pigs significantly increased the intensity of conflict.

Although these are relatively simple predictors, they provide clear and actionable ways to assess vulnerability and to target prevention measures. In summary, our findings indicate that farms with extensive pasture–forest edges, larger numbers of animals, greater domestic animal diversity, and specifically those with cattle and pigs, are at the highest risk of severe conflict. These conditions heighten vulnerability of attacks from spectacled bears, cougars, and feral dogs - the three species most capable of inflicting catastrophic household losses in a single attack, consequently increasing the vulnerability of native carnivores due to retaliation incentives. Our assessment of predictors for conflict intensity with small carnivores also yielded relevant insights, although these models were less significant than those for large carnivores. This may be due to the broader distribution, larger populations, and higher frequency of small carnivore conflicts, which are less influenced by large-scale landscape variables (McKinney, 2002; Nickel et al, 2020). Consistent with this, conflict intensity with small carnivores was not predicted by the amount of pasture–forest edge within farms.

The only variable that significantly predicted higher intensity of conflict with small carnivores was the presence of poultry, which is logical given that poultry represents a primary and physically accessible target for this group (Narváez et al., 2022). No species-level analyses were conducted for small carnivores, as the theoretical framework and overall conflict intensity results already indicated that large carnivores were the priority for understanding conflict dynamics. The spectacled bear (*Tremarctos ornatus*) and feral dogs were analysed separately due to their ecological importance and cultural relevance in Ecuador, whilst the cougar was added to the species-specific analysis due to being the only other ecologically-relevant large carnivore species in the study area.

Finally, it is worth noting that other types of domestic animals were excluded from the models because they were rare in the study area. Livestock in the region is mainly limited to cattle and poultry, which explains their central role in predicting conflict intensity with large and small carnivores, respectively.

### Future conservation directions and limitations of the present study

Beyond interpreting the results for the Ecuadorian Andes, this study also aims to contribute to conservation practice by suggesting management and policy measures informed by our findings. One promising application is the development of vulnerability or risk maps for HWC. By integrating predictor variables such as Pasture–Forest Extent, livestock abundance, and domestic species composition, these models could help identify farms and landscapes at highest risk, enabling more targeted territorial management. In such cases, farmers could be advised to adopt preventive practices, such as avoiding free-roaming cattle near forest edges or experimenting with deterrents (e.g., olfactory or acoustic repellents) (Ugarte et al., 2024; Masini et al., 2005; Shivik et al., 2011; Saurine et al., 2024). Highly vulnerable areas could also be key regions for the implementation of early warning systems (EWS) for conflict situations (UNDRR, 2017). It is equally important to highlight that several key management variables, such as the frequency of human or guard dog presence, and the use of protective measures like electric fencing and secure night enclosures (van Eeden et al., 2018; Petridou et al., 2023; Pimenta et al., 2017; Van Der Weyde et al., 2020), were not assessed in this study. This omission stems from the homogeneity of current livestock management in rural Ecuador, where free-ranging systems without fencing or enclosures dominate (García-Rangel, 2012; MAE, 2019). As a result, there was insufficient variation to test their effects statistically. Nevertheless, evidence from other contexts strongly suggests that such practices substantially reduce predation risk.

Future research should therefore aim to collect larger and more diverse samples that include farms with contrasting management regimes. This would enable robust comparisons and clearer assessments of how specific practices mitigate conflict intensity. Ultimately, supporting the adoption of improved management measures through targeted funding and policy initiatives (for example, investing in the construction of night enclosures for livestock) could help mitigate conflict, safeguard rural livelihoods, and reduce the pressure on vulnerable native carnivores. Lastly, the rising presence of feral dogs and the damage they cause to both native species and livestock present a particularly challenging conservation problem. Any proposed intervention is complicated by the cultural and emotional significance of dogs, which, even in their feral state, are perceived differently from other wildlife. Literature suggests that humans have an evolutionary bond with pets, which significantly reduces antagonism towards them (Figueredo et al., 2023), even when they are causing damage. It is therefore crucial to address this threat without creating further barriers between conservation initiatives and local communities (Slater et al., 2008; Smith et al., 2019). One potential non-lethal measure is the capture and sterilization of feral dogs, which would gradually reduce population growth without resorting to aggressive or lethal control. Extending such programs to unmanaged domestic dog populations could also help prevent the establishment of new feral groups, thereby mitigating both ecological and economic impacts over time. However, it is worth noting that studies on the matter have not yet found strong evidence of successful cases. (Smith et al., 2019).

